# Whole genome and RNA sequencing of 1,220 cancers reveals hundreds of genes deregulated by rearrangement of cis-regulatory elements

**DOI:** 10.1101/099861

**Authors:** Yiqun Zhang, Fengju Chen, Nuno A. Fonseca, Yao He, Masashi Fujita, Hidewaki Nakagawa, Zemin Zhang, Alvis Brazma, Chad J. Creighton, on behalf of the PCAWG Transcriptome Working Group, PCAWG Structural Variation Working Group, ICGC/TCGA Pan-Cancer Analysis of Whole Genomes Network

**Affiliations:** Dan L. Duncan Comprehensive Cancer Center, Baylor College of Medicine, Houston, TX 77030, USA.; Department of Bioinformatics and Computational Biology, The University of Texas MD Anderson Cancer Center, Houston, TX 77030, USA.; Department of Medicine, Baylor College of Medicine, Houston, TX 77030, USA.; Human Genome Sequencing Center, Baylor College of Medicine, Houston, TX 77030, USA.; Laboratory for Genome Sequencing Analysis, RIKEN Center for Integrative Medical Sciences, Tokyo, 108-8639, Japan.; European Molecular Biology Laboratory, European Bioinformatics Institute, EMBL-EBI, Hinxton, UK.; BIOPIC and College of Life Sciences, Peking University, Beijing, 100871, China; Peking-Tsinghua Center for Life Sciences, Peking University, Beijing, 100871, China.

## Abstract

Using a dataset of somatic Structural Variants (SVs) in cancers from 2658 patients—1220 with corresponding gene expression data—we identified hundreds of genes for which the nearby presence (within 100kb) of an SV breakpoint was associated with altered expression. For the vast majority of these genes, expression was increased rather than decreased with corresponding SV event. Well-known up-regulated cancer-associated genes impacted by this phenomenon included *TERT*, *MDM2*, *CDK4*, *ERBB2*, *CD274*, *PDCD1LG2*, and *IGF2*. SVs upstream of *TERT* involved ~3% of cancer cases and were most frequent in liver-biliary, melanoma, sarcoma, stomach, and kidney cancers. SVs associated with up-regulation of PD1 and PDL1 genes involved ~1% of non-amplified cases. For many genes, SVs were significantly associated with either increased numbers or greater proximity of enhancer regulatory elements near the gene. DNA methylation near the gene promoter was often increased with nearby SV breakpoint, which may involve inactivation of repressor elements.

**Abbreviations:**

PCAWG: the Pan-Cancer Analysis of Whole Genomes project
SV: Structural Variant

## Introduction

Functionally relevant DNA alterations in cancer extend well beyond exomic boundaries. One notable example of this involves *TERT*, for which both non-coding somatic point mutations in the promoter or genomic rearrangements in proximity to the gene have been associated with *TERT* up-regulation^1-3^. Genomic rearrangements in cancer are common and often associated with copy number alterations^4, 5^. Breakpoints associated with rearrangement can potentially alter the regulation of nearby genes, e.g. by disrupting specific regulatory elements or by translocating cis-regulatory elements from elsewhere in the genome into close proximity to the gene. Recent examples of rearrangements leading to “enhancer hijacking”— whereby enhancers from elsewhere in the genome are juxtaposed near genes, leading to over-expression—include a distal *GATA2* enhancer being rearranged to ectopically activate *EVI1* in leukemia^6^, activation of GFI1 family oncogenes in medulloblastoma^7^, and 5p15.33 rearrangements in neuroblastoma juxtaposing strong enhancer elements to *TERT*^8^. By integrating somatic copy alterations, gene expression data, and information on topologically associating domains (TADs), a recent pan-cancer study uncovered 18 genes with over-expression resulting from rearrangements of cis-regulatory elements (including enhancer hijacking)^9^. Genomic rearrangement may also disrupt the boundary sites of insulated chromosome neighborhoods, resulting in gene up-regulation^10^.

The Pan-Cancer Analysis of Whole Genomes (PCAWG) initiative has recently assembled over 2600 whole cancer genomes, with high sequencing coverage, from multiple independent studies representing a wide range of cancer types. These data involve a comprehensive and unified identification of somatic substitutions, indels, and structural variants (SVs, representing genomic rearrangement events, each event involving two breakpoints from different genomic coordinates becoming fused together), based on “consensus” calling across three independent algorithmic pipelines, together with initial basic filtering, quality checks, and merging^11,12^. Whole genome sequencing offers much better resolution in SV inference over that of whole exome data or SNP arrays^4,9^. These data represent an opportunity for us to survey this large cohort of cancers for somatic SVs with breakpoints located in proximity to genes. For a sizeable subset of cases in the PCAWG cohort, data from other platforms in addition to whole genome sequencing, such as RNA expression or DNA methylation, are available for integrative analyses, with 1220 cases having both whole genome and RNA sequencing.

While SVs can result in two distant genes being brought together to form fusion gene rearrangements (e.g. *BCR*-*ABL1* or *TMPRSS2*-*ERG*)^13^, this present study focuses on SVs impacting gene regulation in the absence of fusion events or copy number alterations, e.g. SVs with breakpoints occurring upstream or downstream of the gene and involving rearrangement of cis-regulatory elements. With a genome-wide analysis involving a large sample size, information from multiple genes may be leveraged effectively, in order to identify common features involving the observed disrupted regulation of genes impacted by genomic rearrangement.

## Results

### Widespread impact of somatic SVs on gene expression patterns in cancer

Inspired by recent observations in kidney cancer^3,14^, neuroblastoma^8,15^, and B-cell malignancies^16^, of recurrent genomic rearrangements affecting the chromosomal region proximal to *TERT* and resulting in its up-regulation, we sought to carry out a pan-cancer analysis of all coding genes, for ones appearing similarly affected by rearrangement. We referred to a dataset of somatic SVs called for whole cancer genomes of 2658 patients, representing more than 20 different cancer types and compiled and harmonized by the PCAWG initiative from 47 previous studies (Table S1). Gene expression profiles were available for 1220 of the 2658 patients. We set out to systematically look for genes for which the nearby presence of an SV breakpoint could be significantly associated with changes in expression. In addition to the 0-20 kb region upstream of each gene (previously involved with rearrangements near *TERT*^3^), we also considered SV breakpoints occurring 20-50kb upstream of a gene, 50-100kb upstream of a gene, within a gene body, or 0-20kb downstream of a gene (Figure 1a). (SV breakpoints located within a given gene were not included in the other upstream or downstream SV sets for that same gene.) For each of the above SV groups, we assessed each gene for correlation between associated SV event and expression. As each cancer type as a group would have a distinct molecular signature^17^, and as genomic rearrangements may be involved in copy alterations^4^, both of these were factored into our analysis, using linear models.

**Figure 1.**
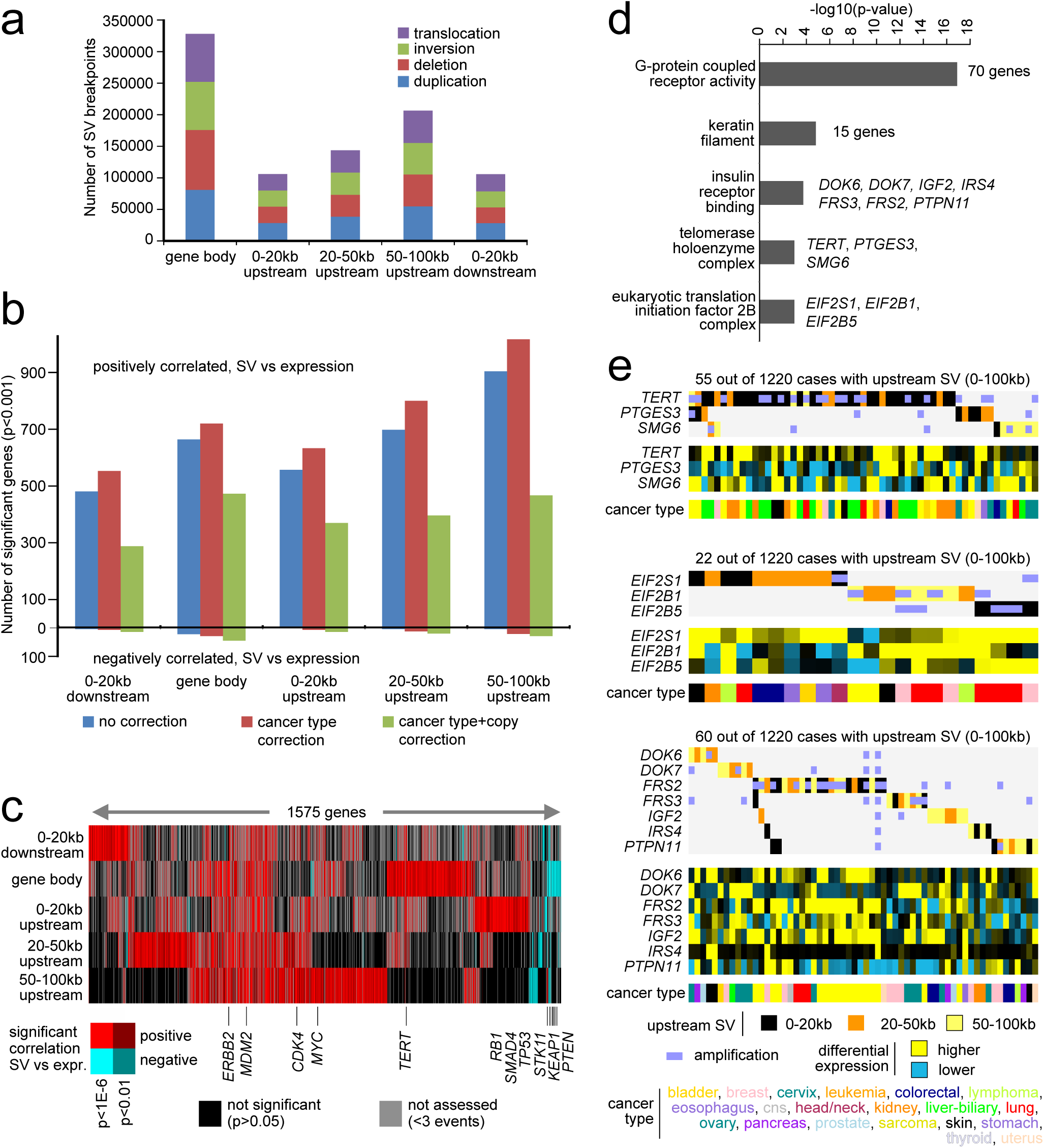
Structural Variants (SVs) associated with altered expression of nearby genes. **(a)** Numbers of SV breakpoints identified as occurring within a gene body, 0-20kb upstream of a gene, 20-50kb upstream of a gene, 50-100kb upstream of a gene, or 0-20kb downstream of a gene. For each SV set, the breakdown by alteration class is indicated. SVs located within a given gene are not included in the other upstream or downstream SV sets for that same gene. **(b)** For each of the SV sets from part a, numbers of significant genes (p<0.001), showing correlation between expression and associated SV event. Numbers above and below zero point of y-axis denote positively and negatively correlated genes, respectively. Linear regression models also evaluated significant associations when correcting for cancer type (red) and for both cancer type and gene copy number (green). **(c)** Heat map of significance patterns for genes from part b (from the model correcting for both cancer type and gene copy number). Red, significant positive correlation; blue, significant negative correlation; black, not significant (p>0.05); gray, not assessed (less than 3 SV events for given gene in the given genomic region). **(d)** Significantly enriched Gene Ontology (GO) terms for genes positively correlated (p<0.001) with occurrence of SV upstream of the gene (for either 0-20kb, 20-50kb, or 50-100kb SV sets). P-values by one-sided Fisher’s exact test. **(e)** Patterns of SV versus expression for selected gene sets from part d (telomerase holoenzyme complex, top; eukaryotic translation initiation factor 2B complex, middle; insulin receptor binding, bottom). Differential gene expression patterns relative to the median across sample profiles. See also Tables S1 and S2 and Supplementary Figure 1.

For each of the genomic regions relative to genes that were considered (i.e. genes with at least three samples associated with an SV breakpoint within the given region), we found widespread associations between SV event and expression, after correcting for expression patterns associated with tumor type or copy number (Figure 1b and Supplementary Figure 1a and Table S2). For gene body, 0-20kb upstream, 20-50kb upstream, 50-100kb upstream, and 0-20kb downstream regions, the numbers of significant genes at p<0.001 (corresponding to estimated false discovery rates^18^ of less than 5%) were 518, 384, 416, 496, and 302, respectively. For each of these gene sets, many more genes were positively correlated with SV event (i.e. expression was higher when SV breakpoint was present) than were negatively correlated (on the order of 95% versus 5%). Permutation testing of the 0-20kb upstream dataset (randomly shuffling the SV event profiles and computing correlations with expression 1000 times) indicated that the vast majority of the significant genes observed using the actual dataset would not be explainable by random chance or multiple testing (with permutation results yielding an average of 30 “significant” genes with standard deviation of 5.5, compared to 384 significant genes found for the actual dataset). Without correcting for copy number, even larger numbers of genes with SVs associated with increased expression were found (Figure 1b), indicating that many of these SVs would be strongly associated with copy gain. Many of the genes found significant for one SV group were also significant for other SV groups (Figure 1c). Tumor purity and total number of SV breakpoints were not found to represent significant confounders (Supplementary Figure 1b). The numbers of statistically significant genes were also found when examining regions further upstream or downstream of genes, up to 1Mb (Supplementary Figure 1c).

### Key driver genes in cancer impacted by nearby SV breakpoints

Genes with increased expression associated with nearby SV breakpoints included many genes with important roles in cancer (Table 1), such as *TERT* (significant with p<0.001 for regions from 0-20kb downstream to 20-50kb upstream of the gene), *MYC* (significant for gene body SV breakpoints), *MDM2* (regions from 0-20kb downstream to 50-100kb upstream), *CDK4* (0-20kb downstream and 20-100kb upstream), *ERBB2* (gene body to 50-100kb upstream), *CD274* (0-20kb downstream to 50-100kb upstream), *PDCD1LG2* (0-20kb downstream to 20-50kb upstream) and *IGF2* (0-20kb downstream and 50-100kb upstream). Genes with decreased expression associated with SV breakpoints located within the gene included *PTEN*^19^ (n=50 cases with an SV breakpoint out of 1220 cases with expression data available), *STK11* (n=15), *KEAP1* (n=5), *TP53* (n=22), *RB1* (n=55), and *SMAD4* (n=18), where genomic rearrangement would presumably have a role in disrupting important tumor suppressors; for other genes, SV breakpoints within the gene could potentially impact intronic regulatory elements, or could represent potential fusion events (though in a small fraction of cases)^13^. Examining the set of genes positively correlated (p<0.001) with occurrence of SV breakpoint upstream of the gene (for either 0-20kb, 20-50kb, or 50-100kb SV sets), enriched gene categories (Figure 1d) included G-protein coupled receptor activity (70 genes), telomerase holoenzyme complex (*TERT*, *PTGES3*, *SMG6*), eukaryotic translation initiation factor 2B complex (*EIF2S1*, *EIF2B1*, *EIF2B5*), keratin filament (15 genes), and insulin receptor binding (*DOK6*, *DOK7*, *IGF2*, *IRS4*, *FRS2*, *FRS3*, *PTPN11*). When taken together, SVs involving the above categories of genes would potentially impact a substantial fraction of cancer cases, e.g. on the order of 2-5% of cases across various types (Figure 1e). Gene amplification events (defined as five or more copies) could be observed for a number of genes associated with SVs, but amplification alone in many cases would not account for the elevated gene expression patterns observed (Figure 1e).

**Table 1.** Genes positively correlated in expression (p<0.001, corrected for copy number and cancer type) with occurrence of upstream SV, with the gene being previously associated with cancer.

| *Region:* | *0-20kb upstream* |  | *20-50kb upstream* |  | *50-100kb upstream* |  | *gene body* |  | *0-20kb downstream* |  |
| --- | --- | --- | --- | --- | --- | --- | --- | --- | --- | --- |
| *Gene* | *n* | *t* | *n* | *t* | *n* | *t* | *n* | *t* | *n* | *t* |
| CDK4 | 16 | 2.39 | 27 | 8.67 | 23 | 5.94 | 13 | 1.92 | 21 | 5.93 |
| ERBB2 | 13 | 3.66 | 17 | 7.99 | 34 | 11.87 | 23 | 8.55 | 15 | 2 |
| MDM2 | 17 | 9.5 | 22 | 7.9 | 21 | 9.52 | 20 | 8.84 | 19 | 8.35 |
| TERT | 31 | 8.08 | 9 | 2.34 | 8 | 0.73 | 10 | 7.39 | 5 | 6.81 |
| CDK12 | 7 | 0.33 | 14 | 3.78 | 14 | -0.02 | 41 | 2.8 | 11 | 3.01 |
| HMGA2 | 10 | 3.71 | 8 | 4.31 | 15 | 1.71 | 24 | 2.16 | 6 | -0.84 |
| EGFR | 8 | 1.69 | 12 | 4.39 | 9 | 2.41 | 31 | 5.57 | 6 | 4.01 |
| TBL1XR1 | 3 | 0.38 | 9 | 3.51 | 9 | 1.11 | 32 | 2.23 | 4 | 2.02 |
| MYCL | 4 | 2.23 | 5 | -0.14 | 10 | 4.24 | 0 | NA | 5 | 3.05 |
| CCND3 | 3 | 2.97 | 6 | 4.01 | 7 | 4.18 | 15 | 4.53 | 5 | 1.44 |
| CLTC | 7 | 1.99 | 4 | 1.66 | 5 | 3.98 | 14 | 0.43 | 6 | 2.93 |
| PDCD1LG2 | 3 | 3.8 | 8 | 4.02 | 4 | 0.97 | 9 | 7.81 | 6 | 5.33 |
| PTPN11 | 4 | 2.83 | 3 | 3.88 | 7 | 2.59 | 7 | 1.1 | 3 | -0.61 |
| SMARCE1 | 2 | NA | 6 | 4.7 | 6 | 3.29 | 6 | 0.75 | 1 | NA |
| PDGFRA | 3 | 3.81 | 4 | 0.07 | 6 | 0.04 | 7 | 1.51 | 2 | NA |
| NF1 | 1 | NA | 3 | 4.44 | 8 | 2.87 | 65 | -2.98 | 0 | NA |
| CD274 | 3 | 3.33 | 3 | 1.64 | 6 | 1.42 | 6 | 5.27 | 4 | 5.1 |
| PRKAR1A | 2 | NA | 3 | 1.3 | 3 | 3.39 | 4 | 2.29 | 1 | NA |
| MYB | 5 | -0.18 | 3 | 3.58 | 0 | NA | 1 | NA | 1 | NA |
| FOXL2 | 2 | NA | 3 | 5.27 | 3 | -0.48 | 0 | NA | 2 | NA |
| BCL7A | 3 | 2.54 | 1 | NA | 3 | 3.38 | 7 | 1.76 | 3 | 2.86 |
| SS18 | 0 | NA | 3 | 3.57 | 4 | 0.49 | 8 | 3.57 | 1 | NA |
| TFE3 | 3 | 3.45 | 1 | NA | 2 | NA | 2 | NA | 0 | NA |
| NKX2-1 | 1 | NA | 3 | 4.24 | 2 | NA | 0 | NA | 1 | NA |

Translocations involving the region 0-100kb upstream of *TERT* were both inter- and intrachromosomal (Figure 2a and Table S3) and included 170 SV breakpoints and 84 cancer cases, with the most represented cancer types including liver-biliary (n=29 cases), melanoma (n=17 cases), sarcoma (n=15 cases), and kidney (n=9 cases). Most of these SV breakpoints were found within 20kb of the *TERT* start site (Figure 2b), which represented the region where correlation between SV events and *TERT* expression was strongest (Figures 2c and 2d, p<1E14, linear regression model). In neuroblastoma, translocation of enhancer regulatory elements near the promoter was previously associated with *TERT* up-regulation^8,15^. Here, in a global analysis, we examined the number of enhancer elements^20^ within a 0.5 Mb region upstream of each rearrangement breakpoint occurring in proximity to *TERT* (for breakpoints where the breakpoint mate was oriented away from *TERT*). While for unaltered *TERT*, 21 enhancer elements are located 0.5 Mb upstream of the gene, on the order of 30 enhancer elements on average were within the 0.5 Mb region adjacent to the *TERT* SV breakpoint (Figure 2e), representing a significant increase (p<1E-6, paired t-test). A trend was also observed, by which SV breakpoints closer to the *TERT* start site were associated with a larger number of enhancer elements (Figure 2d, p<0.03, Spearman’s correlation).

**Figure 2.**
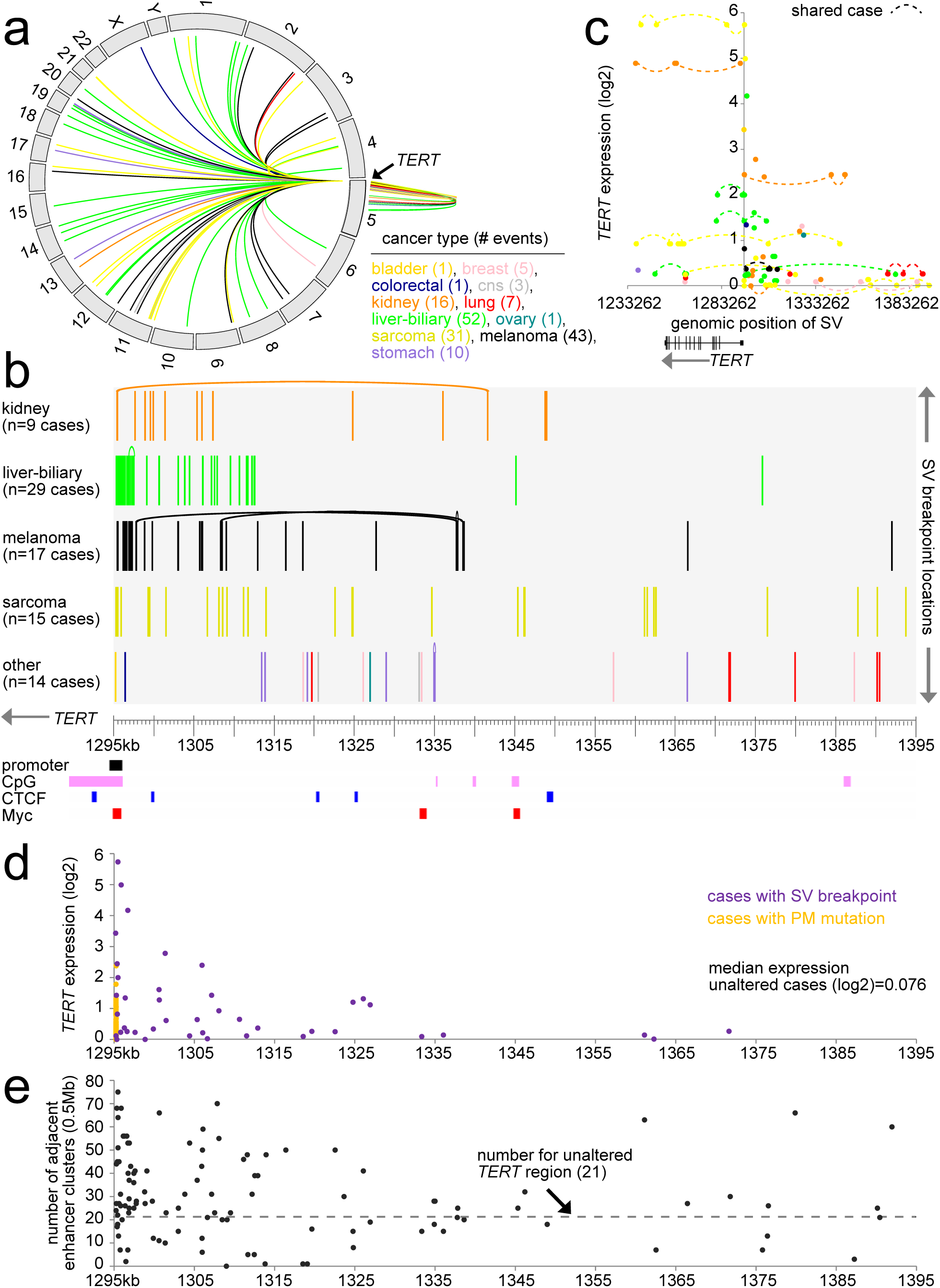
SVs associated with *TERT* and its increased expression. **(a)** Circos plot showing all intra- and interchromosomal rearrangements 0-100kb from the TERT locus. **(b)** By cancer type, SV breakpoint locations within the region ~100kb upstream of *TERT*. Curved line connects two breakpoints common to the same SV. *TERT* promoter, CpG Islands, and CTCF and Myc binding sites along the same region are also indicated. **(c)** Gene expression levels of *TERT* corresponding to SVs located in the genomic region 0-20kb downstream to 100kb upstream of the gene (116 SV breakpoints involving 47 cases). **(d)** Where data available, gene expression levels of *TERT* corresponding to SVs from part b. Expression levels associated with *TERT* promoter (PM) mutation are also represented. Median expression for unaltered cases represents cases without *TERT* alteration (SV, mutation, amplification, viral integration) or *MYC* amplification. For part d, where multiple SVs were found in the same tumor, the SV breakpoint that was closest to the *TERT* start site was used for plotting the expression. **(e)** Numbers of enhancer elements within a 0.5 Mb region upstream of each rearrangement breakpoint are positioned according to breakpoint location. For unaltered *TERT*, 21 enhancer elements were 0.5 Mb upstream of the gene. See also Table S3.

Consistent with observations elsewhere^4^, genomic rearrangements could be associated here with copy alterations for a large number of genes (Figure 1b), including genes of particular interest such as *TERT* and *MDM2* (Figure 3a). However, copy alteration alone would not account for all observed cases of increased expression in conjunction with SV event. For example, with a number of key genes (including *TERT*, *MDM2*, *ERBB2*, *CDK4*), when all amplified cases (i.e. with five or more gene copies) were grouped into a single category, regardless of SV breakpoint occurrence, the remaining SV-involved cases showed significantly increased expression (Figure 3b). Regarding *TERT* in particular, a number of types of genomic alteration may act upon transcription, including upstream SV breakpoint, *TERT* amplification^21^, promoter mutations^1,2^, promoter viral integration^22^, and *MYC* amplification^23^. Within the PCAWG cohort of 2658 cancer cases, 933 (35%) were altered according to at least one of the above alteration classes, with each class being associated with increased *TERT* mRNA expression (Figure 3c). Upstream SVs in particular were associated with higher *TERT* as compared to promoter mutation or amplification events.

**Figure 3.**
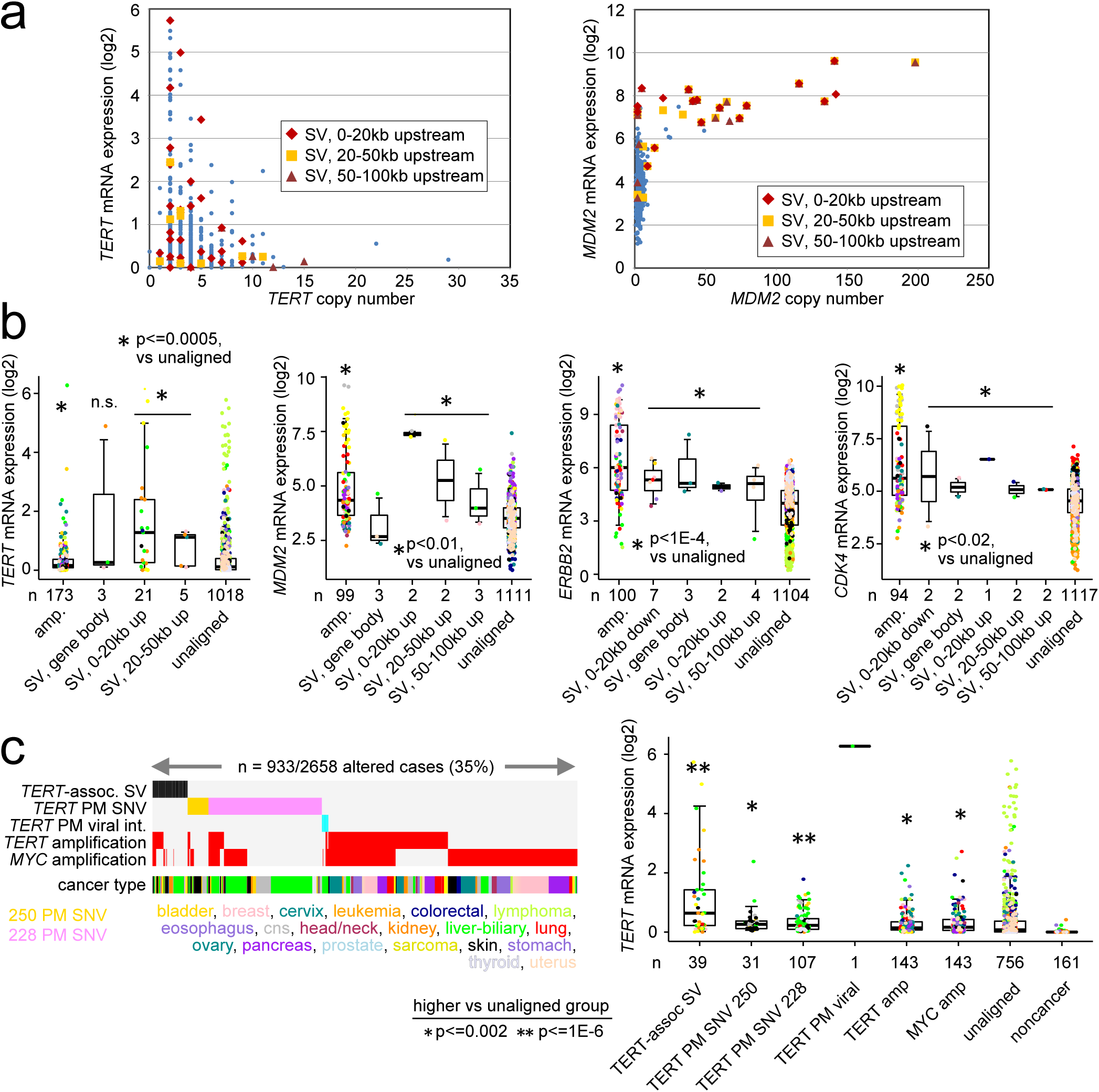
In addition to gene amplification and other genomic alteration events, SVs in proximity to key genes contribute to cases of high expression. **(a)** For 1220 cancer cases, copy number versus expression for *TERT* (left) and *MDM2* (right). Cases with SV events upstream of the gene are indicated. **(b)** Box plots of expression for *TERT*, *MDM2*, *ERBB2*, and *CDK4* by alteration class (“amp.” or gene amplification: 5 or more copies, SV within gene body, SV 0-20kb downstream of gene, SV 0-20kb upstream of gene, SV 20-50kb upstream of gene, SV 50-100kb upstream of gene, or none of the above, i.e. “unaligned”). Cases with both SV and amplification are assigned here within the amplification group. **(c)** Left: Alterations involving *TERT* (SV 0-50kb upstream of gene, somatic mutation in promoter, viral integration within *TERT* promoter, 5 or more gene copies of *TERT* or *MYC*) found in the set of 1220 cancers cases having both WGS and RNA data available. Right: Box plot of *TERT* expression by alteration class. “TERT amp” group does not include cases with other *TERT*-related alterations (SV, Single Nucleotide Variant or “SNV”, viral). P-values by Mann-Whitney U-test. n.s., not significant (p>0.05). Box plots represent 5%, 25%, 50%, 75%, and 95%. Points in box plots are colored according to tumor type as indicated in part c.

SVs associated with *CD274* (PD1) and *PDCD1LG2* (PDL1)—genes with important roles in the immune checkpoint pathway—were associated with increased expression of these genes (Figure 4a and Table S4). Out of the 1220 cases with gene expression data, 19 harbored an SV breakpoint in the region involving the two genes, both of which reside on chromosome 9 in proximity to each other (Figure 4b, considering the region 50kb upstream of *CD274* to 20kb downstream of PDCD1LG2). These 19 cases included lymphoma (n=5), lung (4), breast (2), head and neck (2), stomach (2), colorectal (1), and sarcoma (1). Six of the 19 cases had amplification of one or both genes, though on average cases with associated SV had higher expression than cases with amplification but no SV breakpoint (Figure 4a, p<0.0001 t-test on log-transformed data). For most of the 19 cases, the SV breakpoint was located within the boundaries of one of the genes (Figure 4a), while both genes tended to be elevated together regardless of the SV breakpoint position (Figure 4b). We examined the 19 cases with associated SVs for fusions involving either *CD274* or *PDCD1LG2*, and we identified a putative fusion transcript for *RNF38*->*PDCD1LG2* involving three cases, all of which were lymphoma. No fusions were identified involving *CD274*.

**Figure 4.**
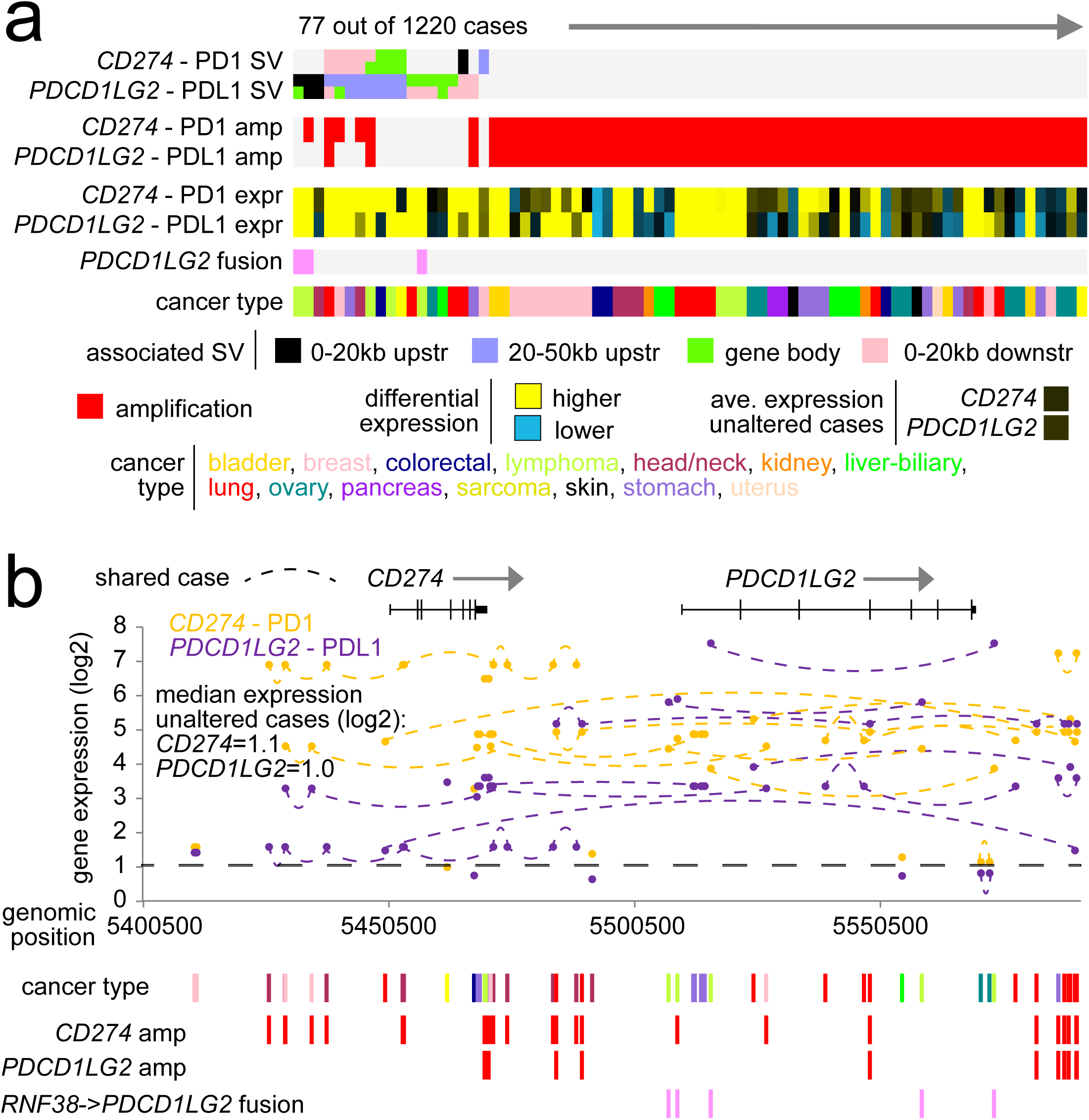
SVs associated with PD1/PDL1 genes and their increased expression. **(a)** Patterns of SV, gene amplification (5 or more copies), *RNF38*->*PDCD1LG2* gene fusion, and differential expression for *CD274* (PD1 gene) and (PDL1 gene), for the subset of cases with associated SV or amplification for either gene. Differential gene expression patterns relative to the median across sample profiles. **(b)** Gene expression levels of *CD274* and of *PDCD1LG2*, corresponding to the position of SV breakpoints located in the surrounding genomic region on chromosome 9 (representing 66 SV breakpoints involving 19 cases). Median expression for unaltered cases represents cases without SV or amplification. See also Table S4.

### SVs associated with translocated enhancers and altered DNA methylation near genes

Similar to analyses focusing on *TERT* (Figure 2d), we examined SVs involving other genes for potential translocation of enhancer elements. For example, like *TERT*, SVs with breakpoints 0-20kb upstream of *CDK4* were associated with an increased number of upstream enhancer elements as compared to that of the unaltered gene (Figure 5a); however, SV breakpoints upstream of *MDM2* were associated with significantly fewer enhancer elements compared to that of the unaltered region (Figure 5a). For the set of 1233 genes with at least 7 SV breakpoints 0-20kb upstream and with breakpoint mate on the distal side from the gene, the numbers of enhancer elements 0.5 Mb region upstream of rearrangement breakpoints was compared with the number for the unaltered gene (Figure 5b and Table S5). Of these genes, 24% showed differences at a significance level of p<0.01 (paired t-test, with ~12 nominally significant genes being expected by chance). However, for most of these genes, the numbers of enhancer elements was decreased on average with the SV breakpoint rather than increased (195 versus 103 genes, respectively), indicating that translocation of greater numbers of enhancers might help explain the observed upregulation for some but not all genes. For other genes (e.g. downstream of *PDCD1LG2 HOXA13* and *CCNE1*), enhancer elements on average were positioned in closer proximity to the gene as a result of the genomic rearrangement (Figure 5c). Of 829 genes examined (with at least 5 SV breakpoints 0-20kb upstream and with breakpoint mate on the distal side from the gene, where the breakpoint occurs between the gene start site and its nearest enhancer in the unaltered scenario), 8.3% showed a significant decrease (p<0.01, paired t-test) in distance to the closest enhancer on average as a result of the SV breakpoint, as compared to 1% showing a significance increase in distance.

**Figure 5.**
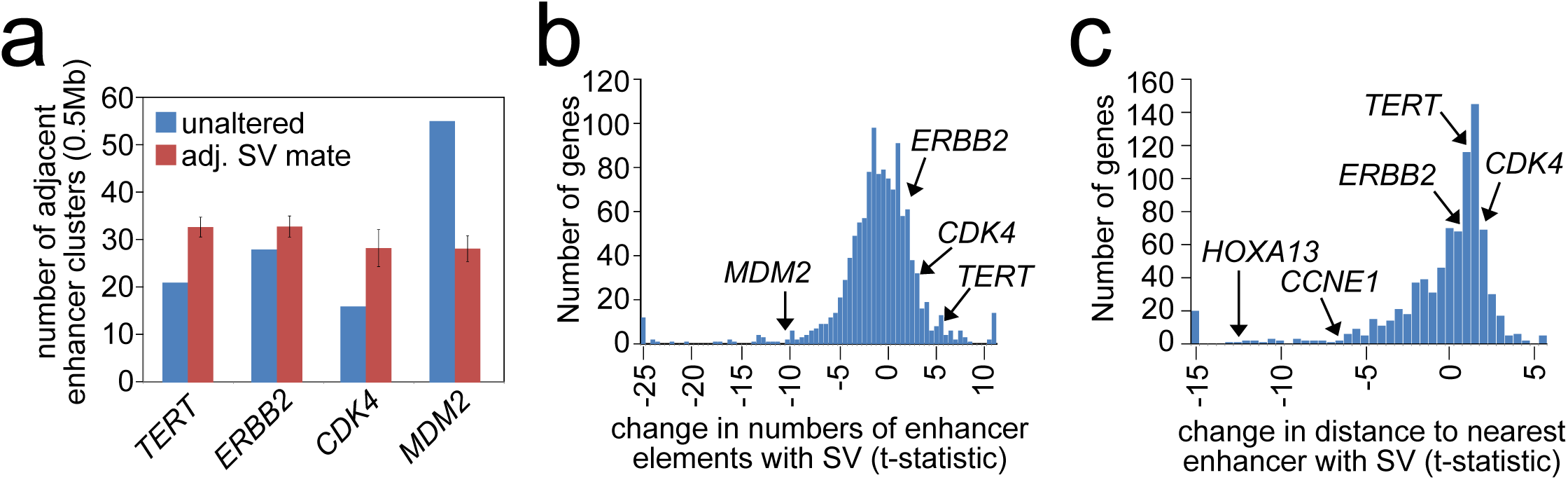
Translocation of enhancer elements associated with SVs near genes. **(a)** For *TERT*, *ERBB2*, *CDK4*, and *MDM2*, average number of enhancer elements within a 0.5 Mb region upstream of each rearrangement breakpoint (considering the respective SV sets occurring 0-20kb upstream of each gene), as compared to the number of enhancers for the unaltered gene. All differences are significant with p<0.01 (paired t-test). Error bars denote standard error. **(b)** For 1233 genes with at least 7 SVs 0-20kb upstream and with breakpoint mate on the distal side from the gene, histogram of t-statistics (paired t-test) comparing numbers of enhancer elements 0.5 Mb region upstream of rearrangement breakpoints with the number for the unaltered gene. Positive versus negative t-statistics denote greater versus fewer enhancers, respectively, associated with the SVs. **(c)** For 829 genes (with at least 5 SVs 0-20kb upstream and with breakpoint mate on the distal side from the gene, where the breakpoint occurs between the gene start site and its nearest enhancer in the unaltered scenario), histogram of t-statistics (paired t-test) comparing the distance of the closest enhancer element upstream of rearrangement breakpoints with the distance for the unaltered gene. Negative t-tatistics denote a shorter distance associated with the SVs. See also Table S5.

We went on to examine genes impacted by nearby SV breakpoints for associated patterns of DNA methylation. Taking the entire set of 8256 genes with associated CpG island probes represented on the 27K DNA methylation array platform (available for samples from The Cancer Genome Atlas), the expected overall trend^24^ of inverse correlations between DNA methylation and gene expression were observed (Figure 6a and Table S6). However, for the subset of 263 genes positively correlated in expression with occurrence of upstream SV breakpoint (p<0.001, 0-20kb, 20-50kb, or 50-100kb SV sets), the methylation-expression correlations were less skewed towards negative (p=0.0001 by t-test, comparing the two sets of correlation distributions in Figure 5a). Genes positively correlated between expression and methylation included *TERT* and *MDM2*, with many of the same genes also showing a positive correlation between DNA methylation and nearby SV breakpoint (Figure 6a). Regarding *TERT*, a CpG site located in close proximity to its core promotor is known to contain a repressive element^8,25^; non-methylation results in the opening of CTCF binding sites and the transcriptional repression of *TERT*^25^. In the PCAWG cohort, SV breakpoints occurring 0-20kb upstream of the gene were associated with increased CpG island methylation (Figure 6b), while SV breakpoints 20-50kb upstream were not; *TERT* promoter mutation was also associated with increased methylation (Figure 6c).

**Figure 6.**
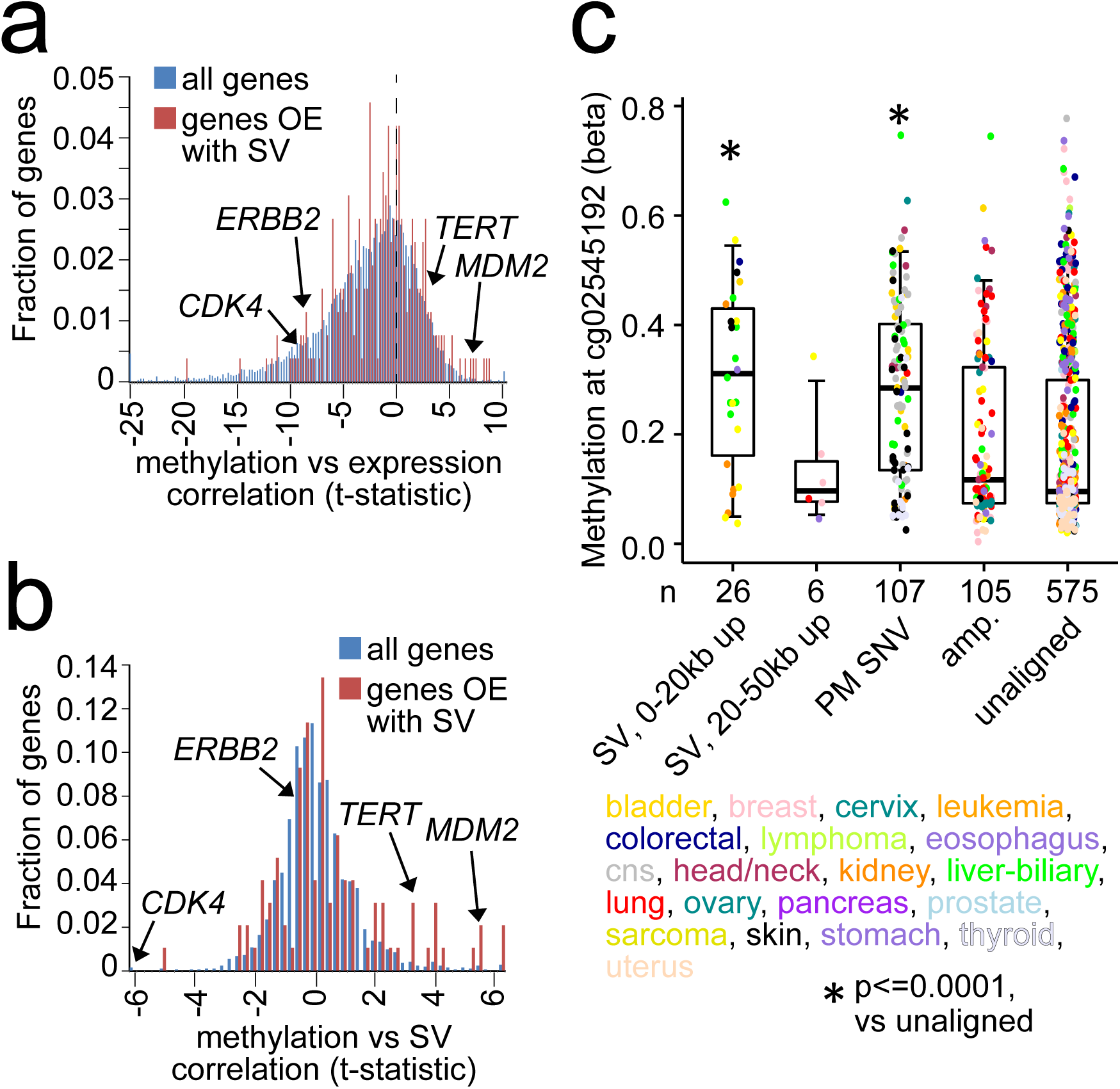
Altered DNA methylation patterns associated with SVs near genes. **(a)** Histogram of t-statistics for correlation between gene expression and DNA methylation (by Pearson’s using log-transformed expression and logit-transformed methylation), for both the entire set of 8256 genes (blue) associated with CpG islands represented on DNA methylation array platform and the subset of 263 genes (red) on methylation platform and positively correlated in expression (p<0.001, “OE” for “over-expressed”) with occurrence of upstream SV (for either 0-20kb, 20-50kb, or 50-100kb SV sets). **(b)** Histogram of t-statistics for correlation between gene expression and SV event (by Pearson’s using logit-transformed methylation), for both the entire set of 2316 genes (blue) with at least 3 SVs 0-20kb upstream and represented on methylation platform and the subset of 97 genes (red) on methylation platform and positively correlated in expression (p<0.001) with occurrence of SV 0-20kb upstream. **(c)** DNA methylation of the CpG site cg02545192 proximal to the TERT core promoter in cases with SV 0-20kb or 20-50kb upstream of TERT, in cases with TERT promoter (PM) activation mutation (SNV), in cases with TERT amplification (“amp.”), and in the rest of cases (unaligned). P-values by t-test on logit-transformed methylation beta values. Box plots represent 5%, 25%, 50%, 75%, and 95%. Points in box plots are colored according to tumor type as indicated. See also Table S6.

## Discussion

Using a unique dataset of high coverage whole genome sequencing and gene expression on tumors from a large number of patients and involving a wide range of cancer types, we have shown here how genomic rearrangement of regions nearby genes, leading to gene up-regulation—a phenomenon previously observed for individual genes such as *TERT*—globally impacts a large proportion of genes and of cancer cases. Genomic rearrangements involved with up-regulation of *TERT* in particular have furthermore been shown here to involve a wide range of cancer types, expanded from previous observations made in individual cancer types such as kidney chromophobe and neuroblastoma. While many of the genes impacted by genomic rearrangement in this present study likely represent passengers rather than drivers of the disease, many other genes with canonically established roles in cancer would be impacted. Though any given gene may not be impacted in this way in a large percentage of cancer cases (the more frequently SV-altered gene *TERT* involving less than 3% of cancers surveyed), the multiple genes involved leads to a large cumulative effect in terms of absolute numbers of patients. The impact of somatic genomic rearrangements on altered cis-regulation should therefore be regarded as an important driver mechanism in cancer, alongside that of somatic point mutations, copy number alteration, epigenetic silencing, gene fusions, and germline polymorphisms.

While the role of genomic rearrangements in altering the cis-regulation of specific genes within specific cancer types has been previously observed, our present pan-cancer study demonstrates that this phenomenon is more extensive and impacts a far greater number of genes than may have been previously thought. A recent study by Weischenfeldt et al.^9^, utilizing SNP arrays to estimate SV breakpoints occurring within TADs (which confine physical and regulatory interactions between enhancers and their target promoters), uncovered 18 genes (including *TERT* and *IRS4*) in pan-cancer analyses and 98 genes (including *IGF2*) in cancer type-specific analyses with over-expression associated with rearrangements involving nearby or surrounding TADs. Our present study using PCAWG datasets identifies hundreds of genes impacted by SV-altered regulation, far more than the Weischenfeldt study. In contrast to the Weischenfeldt study, our study could take advantage of whole genome sequencing over SNP arrays, with the former allowing for much better resolution in identifying SVs, including those not associated with copy alterations. In addition, while TADs represent very large genomic regions, often extending over 1Mb, our study pinpoints SV with breakpoints acting within relatively close distance to the gene, e.g. within 20kb for many genes. In principle, genomic rearrangements could impact a number of regulatory mechanisms, not necessarily limited to enhancer hijacking, and genes may be altered differently in different samples. The analytical approach of our present study has the advantage of being able to identify robust associations between SVs and expression, without making assumptions as to the specific mechanism.

Future efforts can further explore the mechanisms involved with specific genes deregulated by nearby genomic rearrangements. Regarding *TERT*-associated SVs, for example, previously observed increases in DNA methylation of the affected region had been previously thought to be the result of massive chromatin remodeling brought about by juxtaposition of the *TERT* locus to strong enhancer elements^8^, which is supported by observations made in this present study involving multiple cancer types. However, not all genes found here to be deregulated by SVs would necessarily follow the same patterns as those of *TERT*. For example, not all of the affected genes would have repressor elements being inactivated by DNA methylation, and some genes such as *MDM2* do not show an increase in enhancer numbers with associated SV breakpoints but do correlate positively between expression and methylation. There is likely no single mechanism that would account for all of the affected genes, though some mechanisms may be common to multiple genes. Integration of other types of information (e.g. other genome annotation features, data from other platforms, or results of functional studies) may be combined with whole genome sequencing datasets of cancer, in order to gain further insights into the global impact of non-exomic alterations, where the datasets assembled by PCAWG in particular represent a valuable resource.

## Acknowledgements

This work was supported in part by National Institutes of Health (NIH) grant P30CA125123 (C. Creighton) and Cancer Prevention and Research Institute of Texas (CPRIT) grant RP120713 C2 (C. Creighton).

## Methods

### Datasets

Datasets of structural variants (SVs), RNA expression, somatic mutation, and copy number were generated as part of the Pan-Cancer Analysis of Whole Genomes (PCAWG) project.^11,12^ In all, 2671 patients with whole genome data were represented in the PCAWG datasets, spanning a range of cancer types (bladder, sarcoma, breast, liver-biliary, cervix, leukemia, colorectal, lymphoma, prostate, eosophagus, stomach, central nervous system or “cns”, head/neck, kidney, lung, skin, ovary, pancreas, thyroid, uterus). For SVs, calls were made by three different data centers using different algorithms; calls made by at least two algorithms were used in the downstream analyses, along with additional filtering criteria being used as described elsewhere^11,26^. The consensus SV calls are available from synapse (https://www.synapse.org/#!Synapse:syn7596712).

For copy number, the calls made by the Sanger group were used. Gene copies of five or more were called as amplification events. For somatic mutation of *TERT* promoter, PCAWG variant calls, as well as any additional data available from the previous individual studies^3,22,27,28^, were used. *TERT* promoter viral integrations were obtained from ref^22^. Of the 2658 cases, RNA-seq data were available for 1220 cases. For RNA-seq data, alignments by both STAR and TopHat2 were used to generated a combined set of expression calls; FPKM-UQ values (where UQ= upper quartile of fragment count to protein coding genes) were used (dataset available at https://www.synapse.org/#!Synapse:syn5553991). Where a small number of patients had multiple tumor sample profiles, one profile was randomly selected to represent the patient. DNA methylation profiles had been generated for 771 cases by The Cancer Genome Atlas using either the Illumina Infinium HumanMethylation450 (HM450) or HumanMethylation27 (HM27) BeadChips (Illumina, San Diego, CA), as previously described^29^. To help correct for batch effects between methylation data platforms (HM450 versus HM27), we used the combat software^30^. For each of 8226 represented genes, an associated methylation array probe mapping to a CpG island was assigned; where multiple probes referred to the same gene, the probe with the highest variation across samples was selected for analysis. Correlations between DNA methylation and gene expression were assessed using logit-transformed methylation data and log-transformed expression data and Pearson’s correlations.

### Integrative analyses between SVs and gene expression

For each of a number of specified genomic region windows in relation to genes, we constructed a somatic SV breakpoint matrix by annotating for every sample the presence or absence of at least one SV breakpoint within the given region. For the set of SVs associated with a given gene within a specified region in proximity to the gene (e.g. 0-20kb upstream, 20-50kb upstream, 50-100kb upstream, 0-20kb downstream, or within the gene body), correlation between expression of the gene and the presence of at least one SV breakpoint was assessed using a linear regression model (with log-transformed expression values). In addition to modeling expression as a function of SV event, models incorporating cancer type (one of the 20 major types listed above) as a factor in addition to SV, and models incorporating both cancer type and copy number in addition to SV, were also considered. For these linear regression models, genes with at least three samples associated with an SV breakpoint within the given region were considered. Genes for which SVs were significant (p<0.001) after correcting for both cancer type and copy were explored in downstream analyses. Results from both the SV only model and results from the SV+cancer type models were also highlighted in Figure 1 and provided in Table S2, but the p-values from those models were not used in selecting for genes or SVs of interest for follow-up analyses.

The method of Storey and Tibshirani^18^ was used to estimate false discovery rates (FDR) for significant genes. For purposes of FDR, only genes that had SV breakpoints falling within the given region relative to the gene in at least 3 cases were tested; for example, for the 0-20kb upstream region, 6257 genes were tested, where 384 genes were significant at a nominal p-value of <'0.001 (using a stringent cutoff, with ~6 genes expected by chance due to multiple testing, or FDR<2%); the other genomic region windows yielded similar results. For each genomic region window, the FDR for genes significant at the p<0.001 level did not exceed 4%. In addition, permutation testing of the 0-20kb upstream dataset was carried out, whereby the SV events were randomly shuffled (by shuffling the patient ids) and the linear regression models (incorporating both cancer type and copy number) were used to compute expression versus permuted SV breakpoint associations; for each of 1000 permutation tests, the number of nominally significant genes at p<0.001 was computed and compared with results from the actual datasets.

### Integrative analyses using enhancer genomic coordinates

Gene boundaries and locations of enhancer elements were obtained from Ensembl (GRCh37 build). Enhancer elements found in multiple cell types (using Ensembl “Multicell” filter, accessed April 1, 2016) were used^20^. As previously described^20^, the Ensembl team first reduced all available experimental data for each cell type into a cell type-specific annotation of the genome; consensus “Multicell” regulatory features of interest, including predicted enhancers, were then defined. For each SV breakpoint 0-20kb upstream of a gene, the number of enhancer elements near the gene that would be represented by the rearrangement was determined (based on the orientation of the SV breakpoint mate). Only SVs with breakpoints on the distal side from the gene were considered in this analysis; in other words, for genes on the negative strand, the upstream sequence of the breakpoint should be fused relative to the breakpoint coordinates, and for genes on the positive strand, the downstream sequence of the breakpoint (denoted as negative orientation) should be fused relative to the breakpoint coordinates.

### Statistical Analysis

All P-values were two-sided unless otherwise specified.

## Figure Legends

**Supplementary Figure 1.**
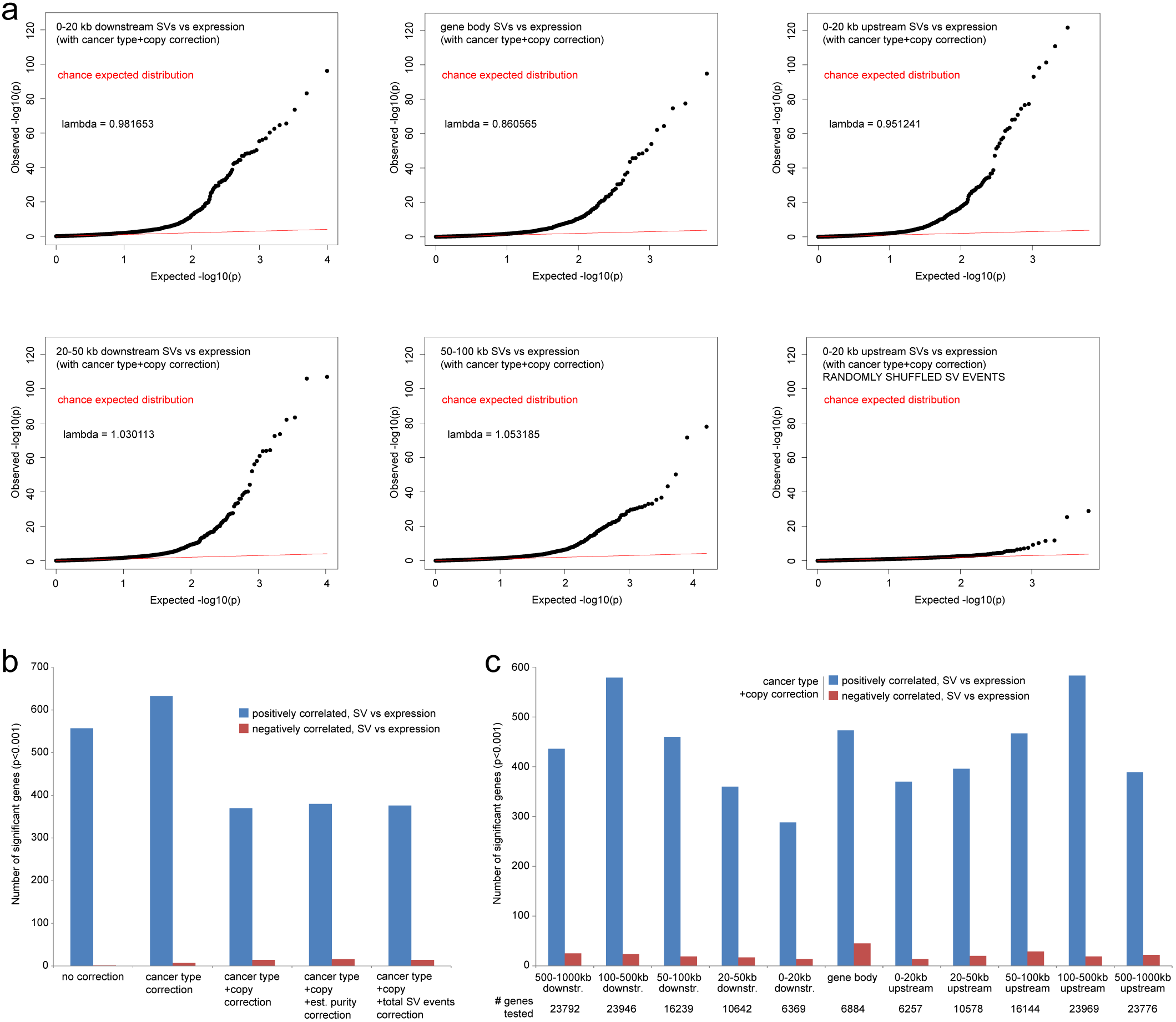
Additional analyses involving the identifications of SVs associated with altered expression of nearby genes. **(a)** QQ plots of the linear regression p-values from the SV events versus expression correlations (correcting for both cancer type and copy number), for SV breakpoints falling within each of the indicated regions surrounding the gene. The bottom right QQ plot represents results from the 0-20kb upstream SV events being randomly permuted relative to the expression profiles. R code for generating q-q plots publicly available from Dr. Stephen Turner (http://www.gettinggeneticsdone.com/2010/07/qq-plots-of-p-values-in-r-using-base.html). Lamda values represent [median of p-values]/0.456, intended as a measure of the observed versus expected median (the concept being borrowed from Genome-Wide Association Studies and applied here to the differential expression analyses). **(b)** Numbers of significant genes (p<0.001, linear regression model), showing correlation between expression and associated SV events (for breakpoints occurring 0-20kb upstream of the gene). Linear regression models represented evaluated significant associations (1) without corrections for other factors, (2) when correcting for cancer type, (3) when correcting for both cancer type and gene copy number, (4) when correcting for cancer type and gene copy number and estimated tumor purity, and (5) when correcting for cancer type and gene copy number and total number of sample-level gene-associated SV events (0-20kb upstream). **(c)** Numbers of significant genes showing correlation between expression and associated SV event (p<0.001, linear regression model incorporating both cancer type and gene copy number), for SVs occurring 500-1000kb downstream of the gene, 100-500kb downstream of the gene, 50-100kb downstream of the gene, 20-50kb downstream of the gene, 0-20kb downstream of the gene, within the gene body, 0-20kb upstream of the gene, 20-50kb upstream of the gene, 50-100kb upstream of the gene, 100-500kb upstream of the gene, and 500-1000kb upstream of the gene.

## Supplementary Data Files

**Table S1, related to Figure 1.** Cancer cases examined in this study, with patient-level annotation regarding specific molecular features.

**Table S2, related to Figure 1.** Complete set of correlations between gene expression and nearby SV event, according to region examined and the regression model applied.

**Table S3, related to Figure 2.** SVs associated with *TERT*, with associated expression and numbers of enhancer elements within a 0.5 Mb region upstream of each rearrangement breakpoint.

**Table S4, related to Figure 4.** SVs associated with *CD274* and *PDCD1LG2*, with associated expression.

**Table S5, related to Figure 5.** Includes the following: 1) Numbers of enhancer elements within a 0.5 Mb region upstream of each rearrangement breakpoint, with associated enrichment patterns (for 1233 genes with at least 7 SVs 0-20kb upstream and with breakpoint mate on the distal side from the gene); and 2) Average change in distance of the nearest enhancer element in proximity to each gene, as a result of rearrangement (for 829 genes with at least 5 SVs 0-20kb upstream and with breakpoint mate on the distal side from the gene, where the breakpoint occurs between the gene start site and its nearest enhancer in the unaltered scenario).

**Table S6, related to Figure 6.** Correlations between DNA methylation and gene expression, and correlations between DNA methylation and adjacent SV event (for breakpoints occurring 0-20kb upstream).

